# Innovations in skin aging: biomimetic exosomes *versus* natural exosomes

**DOI:** 10.1101/2025.01.03.631233

**Authors:** Antonio Salvaggio, Anna Privitera, Giulia Iannello, Maria Violetta Brundo

## Abstract

In this study, we compared the effectiveness of two commercial products contain biomimetic and natural exosomes on skin aging used two cells line, keratinocytes and fibroblasts. MTT assay and scratch-wound assay were employed to evaluate their *in vitro* effects. The treatment with product containing natural exosomes purified from bovine colostrum and loaded with growth factors and cytokines purified from bovine colostrum (AMPLEX PLUS technology), was able to significantly enhance cell proliferation of fibroblasts and keratinocytes at 24 hours compared to cells treatment with product containing biomimetic exosomes and compared to untreated cells (control). Product containing natural exosomes was also able to close the wound scratch after 24 hours. The obtained results suggest that the product containing exosomes derived from bovine colostrum has excellent structural and functional stability offer great potential as natural product for skin aging damage and wound healing.

## 1. Introduction

Skin aging is a complex phenomenon determined by both intrinsic and extrinsic factors, and is influenced by a variety of elements, including genetic predisposition, epigenetic factors, and environmental influences [1, 2]. As a rule, a distinction is made between biological or chronological aging and photoaging, in which the alterations are largely attributable to the cumulative action of the sun’s rays [3]. Many of the signs commonly attributed to chronological aging are actually due to sun damage [3]. Generally, the skin shows the first signs of aging during the fourth decade of life, and it is reasonable to believe that molecular alterations are the basis of this process [4]. Skin aging is characterized by a decreased rate of cell reproduction and an alteration of all repair and renewal processes [5,6].

The aging of the epidermis must be related in relation to the aging of the dermis [7]. In fact, while epidermal cells show aging due to maturation and migrate outward to form a keratinized layer, the dermis undergoes changes related to the number of cells and elastic content [7]. As age increases, the fibrous part of the dermis increases more than the elastic portion; in fact, with age, there is a reduction in turnover and a reduction in nutritional exchange, such that the dermis can no longer carry out its function normally. The dermis is a fibrous structure, principally composed of collagen, elastin, fibronectin, and proteoglycans [8]. These ECM proteins constitute the major of the dermis, providing the skin with tensile strength and mechanical properties. Dermal fibroblasts synthesize, organize and maintain the collagen-rich ECM in human skin [8]. Skin aging prevention and therapy have been the subject of research for decades. Of particular interest are new techniques through which optimal results can be achieved without side effects [9]. Liposomes are artificial vesicles assembled from phospholipids to form a spherical bilayer structure surrounding a small aqueous area. Liposomes have variable dimensions between 50 and 1000 nm and are classified based on their diameter (ultra-small, small, medium and large) and structure (unilamellar, multilamellar, or multivesicle) [10]. These characteristics significantly affect their plasma half-life and degree of loading. Phospholipids are molecules composed of a hydrophilic head and a hydrophobic tail, and in aqueous solution, they spontaneously themselves in a double layer with the tails inside and, if appropriately stimulated, they can form a stable sphere containing a small part of liquid in which they are immersed. Liposomes have good biocompatibility and biodegradability and effectively isolate the product inside them. Unfortunately, liposomes also present certain limitations, including poor targeting, instability, short *in vivo* circulation time, and low utilization of encapsulated substances [11]. In order to solve these problems, a wide variety of delivery systems have been developed in recent years to enhance therapeutic efficacy and reduce side effects. Nanotechnology has been introduced into medicine to improve these products, and artificial or biomimetic exosomes have been developed to carry different therapeutic factors [12]. Biomimetic exosomes are characterized by a good biocompatibility and potential for large-scale production [13], but compared to natural exosomes, they have different disadvantages [13-17], and their damage could cause is not yet well understood, as sufficient *in vitro* and *in vivo* studies have not been conducted, and there are no clinical studies on their effects.

These problems can be overcome by using natural nanovesicles. Exosomes are natural nanovesicles produced by all known cells and have unique characteristics allowing them to function as highly efficient natural nanocarriers. They have inherent stability, low immunogenicity, biocompatibility and good ability to penetrate biological membranes. Increasingly, studies have shown that exosomes can regulate a variety of biological functions and are a vital source of biomarkers for clinical diagnosis [18].

The aim of the present study is to evaluate the effectiveness of commercial product that use biomimetic or natural exosomes for the reduction of skin aging.

## 2. Materials and Methods

For our research we purchased two commercial products utilizing new technologies containing among the ingredients artificial or natural exosomes. In particular, one product contains 25 billion biomimetic exosomes loaded with sodium and potassium salts, sugars, phospholipids, lipids, oligopeptide and several polypeptides (indicated with Biomimetic-Exo). The second compared product contains exosomes purified from bovine colostrum passively loaded with growth factors and cytokines purified from bovine colostrum (indicated with Colostrum-Exo). This latest technology is called AMPLEX plus technology. This product contains 5 mL of AMPLEX plus that includes 20 billion exosomes and 200 mg of growth factors. Both products contain hyaluronic acid.

### 2.1 Propagation and maintenance of cells

Normal adult human dermal fibroblasts (HDFa) (ThermoFisher Scientific) and normal adult primary human epidermal keratinocytes (HEKa) (Invitrogen) were maintained in a humidified incubator with 5% CO_2_ at 37°C in specific medium (Dulbecco’s Modified Eagle and Epilife Medium respectively). Cells were cultured in supplemented with 10% FBS, streptomycin (0.3 mg mL^−1^) and penicillin (50 IU mL^−1^), and GlutaMAX (1 mM) using 75 cm^2^ polystyrene culture flasks and were split every 2–3 days depending on cell confluence.

### 2.2 Cell proliferation assay

Keratinocytes and fibroblasts were treated with two products at three different concentrations (0.5%, 1% and 2%) and incubated for 24 hours in a humidified environment (37 °C and 5% CO_2_). Cells were incubated in absence or presence of 10% FBS. Cells were incubated with and without 10% FBS. Cells incubated with only 10% FBS were used as negative control. As positive control was treatment with 80 µM H_2_O_2_. At the end of the treatment, MTT solution (1 mg/mL) was added to each well and cells were incubated for 2 hours in a humidified environment (37 °C and 5% CO_2_). Epoch Microplate Spectrophotometer (Bio Tec) was used to read the absorbance at 569 nm. Values were normalized with respect to control untreated cells and were expressed as the percent variation of cell proliferation.

### 2.3 Scratch-wound assay

Both cell lines were plated into six-well plates at the density of 3 x 10^6^ cells/well allowed to incubate until confluence. Once the confluence was reached, cells were scraped by using a 1 mL sterile. Cells were washed by using PBS to remove detached cells before adding the medium, in absence or presence of tested products used at 2%. Untreated cells were considered as control. Evaluation was performed 24 hours after scratching.

### 2.4 Statistical analysis

The statistical analysis was carried out by using Graphpad Prism software (version 8.0) (Graphpad software, San Diego, CA, USA). Two-way analysis of variance (ANOVA), followed by Tukey’s post hoc test, was used for multiple comparisons. Statistical significance was set at p-values < 0.01; very significant for p-value < 0.001; highly significant for p-value < 0.0001. Data were reported as the mean ± SD of at least three independent experiments.

## 3. Results and Discussion

Many studies suggest that exosomes have an important impact on skin regeneration, especially in chronic wounds [19, 20]. This role seems to be due to the action they have on the anti-inflammatory response, in fact exosomes contribute to modulation of cell, regulation of inflammation and immune environment, promotion of collagen deposition and angiogenesis [20, 21]. For their ability to transfer signaling molecules to regulate the physiological state of target cells, exosomes can be used for delivering several factors [22]. Therefore, exosomes represent a promising treatment for wound management [23, 24]. Exosomes are divided into natural and engineered exosomes. Biomimetic exosomes are synthetic particles engineered using basic fractions present in natural exosomes (lipids, RNA, proteins) to reproduce their effects. The aim of our research was to evaluate the difference between two commercial products that use the two types of exosomes, biomimetic and natural. In particular, we evaluated their effectiveness on the maintenance and viability of two normal human cell lines, keratinocytes and fibroblasts, testing three different concentrations with or without 10% of fetal bovine serum. The tests showed that the product containing biomimetic exosomes loaded with sodium and potassium salts, sugars, phospholipids, lipids, oligopeptide and several polypeptides, significantly reduced cell viability in the samples treated only with the product at all three concentrations tested and in both cell lines (Figs 1A-1F). A slight decrease in viability was also found in the samples treated with the three concentrations of biomimetic exosomes plus 10% FBS (Figs 1A-1F).

**Figure 1.**
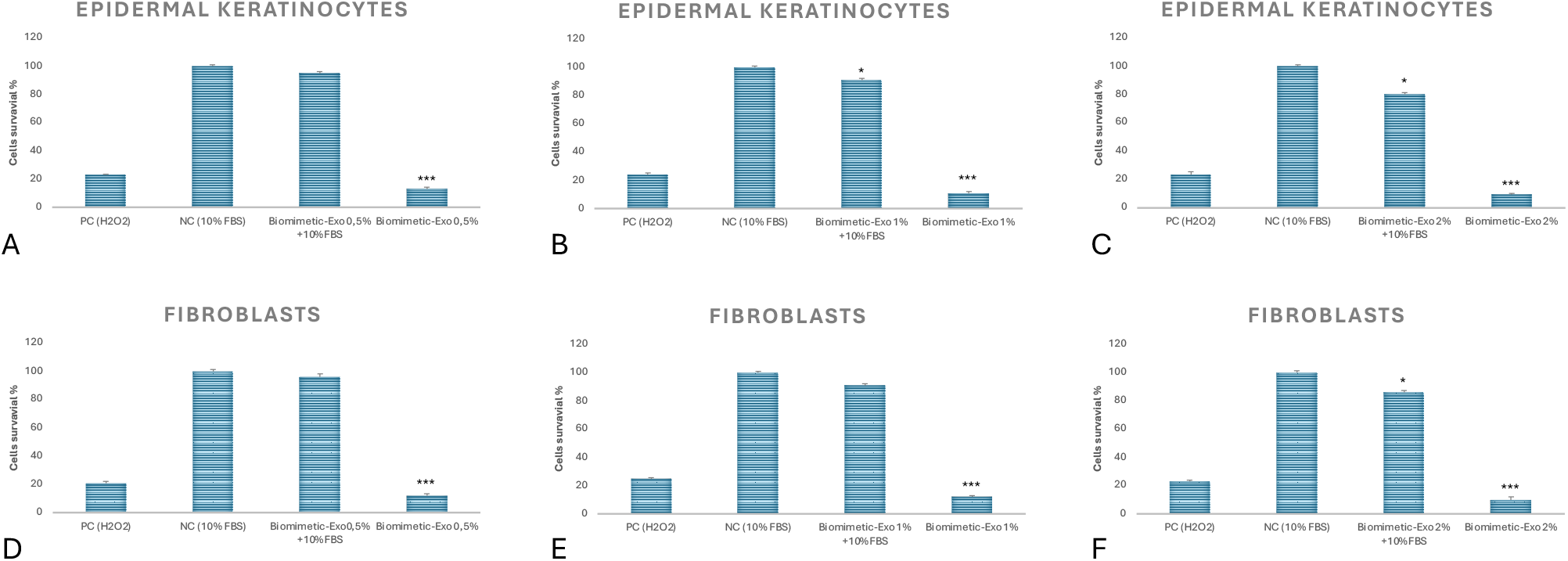
Epidermal keratinocytes (**A**-**C**) and fibroblasts (**D**-**F**) survival following treatment with product contains biomimetic exosomes. Tested three different concentrations: **A** and **D**) 0.5%; **B** and **E**) 1%; **C** and **F**) 2%) with 10% FBS and without 10% FBS. NC (Negative Control, cells, only 10% FBS). PC (Positive Control, 80 µM H_2_O_2_). The asterisks denote the degree of significance between results: ***p<0.0001. Errors bars represent the Standard Deviation of the mean (experiment was repeated three times).

In both cell lines treated with natural exosomes extracted from bovine colostrum and loaded with growth factors and cytokines extracted from colostrum, however, a highly significant difference was found compared to untreated cells (negative control) for all three concentrations. tested when it was used only in products without 10% FBS compared to those treated also with 10% FBS (Figs 2A-2F). These results depend on the fact that osmolarity is modified and therefore cell growth is inhibited. Since we obtained the best results with the 2% concentration, we used this to do the scratch test.

**Figure 2.**
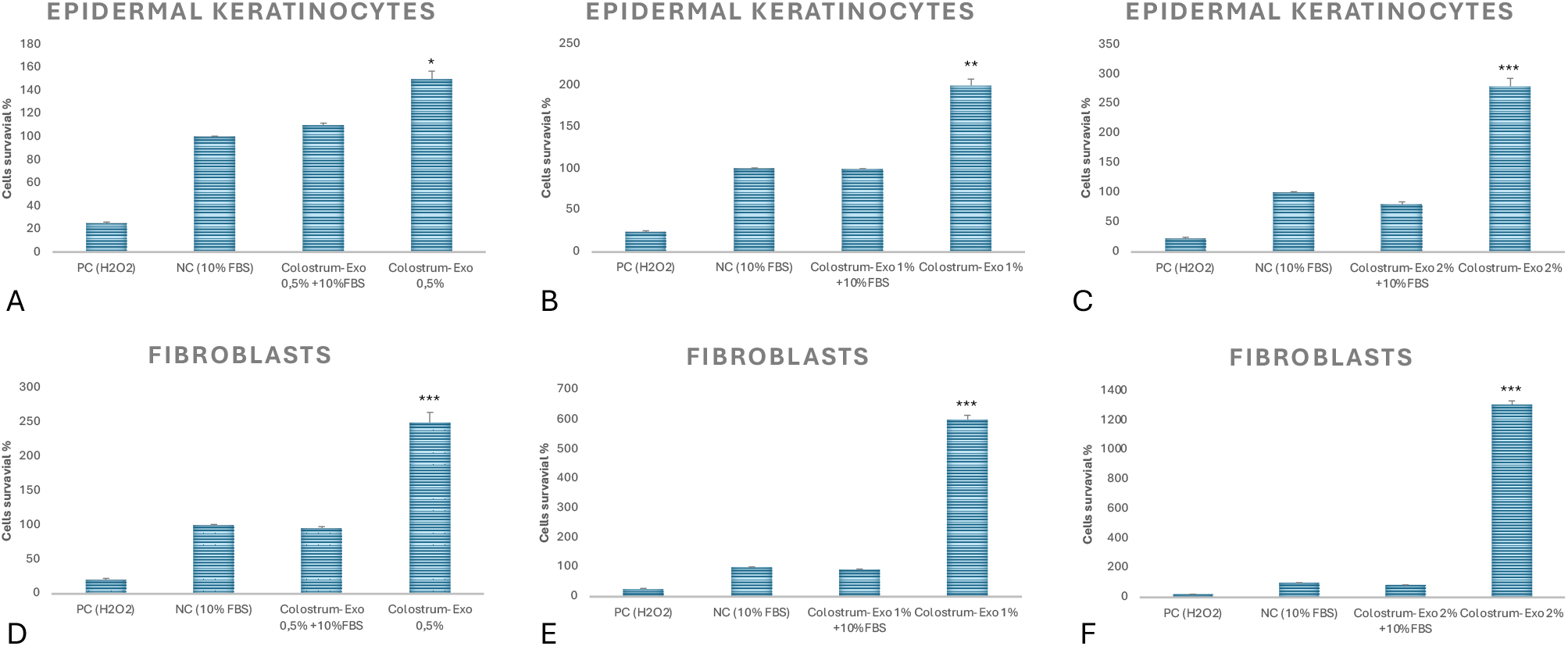
Epidermal keratinocytes (**A**-**C**) and fibroblasts (**D**-**F**) survival following treatment with product contains colostrum exosomes. Tested three different concentrations: **A** and **D**) 0.5%; **B** and **E**) 1%; **C** and **F**) 2%) with 10% FBS and without 10% FBS. NC (Negative Control, cells, only 10% FBS). PC (Positive Control, 80 µM H_2_O_2_). The asterisks denote the degree of significance between results: ***p<0.0001. Errors bars represent the Standard Deviation of the mean (experiment was repeated three times).

**Figure 3.**
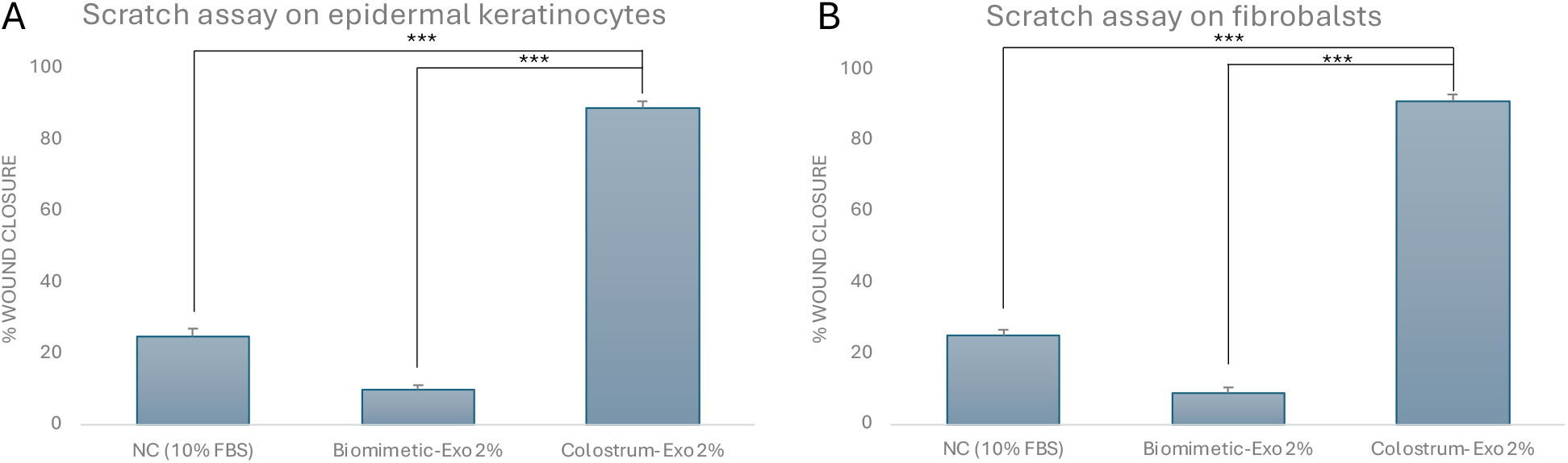
Scratch assay on human epidermal keratinocytes (**A**) and human fibroblasts (**B**) treated with the two products to 2% concentration at 24 hours’ time. Untreated cells were considered as a control (NC). The asterisks denote the degree of significance between results: ***p<0.0001. Errors bars represent the Standard Deviation of the mean (experiment was repeated three times).

Since we obtained the best results with the 2% concentration, we used it to perform the scratch test, which only after 24 hours showed a wound closure capacity of 89% and 91% for epidermal keratinocytes and fibroblasts respectively in samples treated with 2% exosomes extracted and purified from colostrum. The difference compared to cells treated with biomimetic exosomes (10% for keratinocytes and 9% for fibroblasts) and untreated cells (25% for both cell lines) was highly significant (p<0.0001).

The results obtained, certainly, depend both on the factors loaded in the two types of exosomes, but also on their ability to interact with the target molecules present on the plasma membranes of the cells. Colostrum exosomes are loaded with 20 different biologically active factors purified from bovine colostrum, which influence cell survival, differentiation, proliferation, and control over migration, necessary to provide a local environment for tissue regeneration [25]. The factors particularly relevant for tissue healing are TGF-β, IGF-1 and EGF control numerous cellular responses, increase epithelial cell migration and stimulate the collagen secretion by fibroblasts [26, 27]. IGF-1 involved in cell proliferation, differentiation, and individual growth, especially in cartilage and muscle tissue. The bFGF and VEGF are the strongest vascular growth factors involved in the process of proliferation, differentiation, and survival of a wide variety of cell types [26, 27]. Bovine colostrum is an easily accessible, abound source of exosomes, which can be isolated in super-high yields using methods such as ultra-centrifugation [28], overcoming one of the primary limitations of natural exosomes, namely the challenge of producing them on an industrial scale. Recent studies have demonstrated that exosomes purified from bovine colostrum, with their excellent structural and functional stability, can enhance collagen production while reducing reactive oxygen species (ROS) and melanin levels in various cell types. These findings suggest their potential applicability in advanced cell-free skin regeneration therapies [29].

## 4. Conclusions

In recent years, there has been growing interest in the study both natural and engineered exosomes purified from different matrices, especially for their role in wound healing [30, 31]. Exosomes play a crucial role in wound repair and show great promise as important drug delivery vehicles. Exosomes purified from some matrices, such as colostrum, contain specific active components that are directly involved in the wound healing process. Although engineered exosomes could be loaded with specific factors, their membrane phospholipids cannot fully replicate all the properties of natural exosomes, and large-scale production would be impractical [12-17]. This study compared two commercial products using biomimetic and natural exosomes, demonstrating a great effectiveness of natural exosomes compared to biomimetic ones in maintaining cell vitality, promoting wound closure, and supporting overall wound healing.

## Author Contributions

Conceptualization, A.S. and M.V.B.; methodology, A.P. and G.I.; software, A.P.; validation, A.P.; formal analysis, G.I.; data curation, A.S..; writing-original draft preparation, A.S. and M.V.B.; writing-review and editing, A.S., and M.V.B.; supervision, A.S. and M.V.B.; funding acquisition, M.V.B. All authors have read and agreed to the published version of the manuscript.

## Funding

This research received no external funding.

## Institutional Review Board Statement

This study was performed in line with the principles of the Declaration of Helsinki and does not require approval by the Ethics Committee of University of Catania.

## Informed Consent Statement

Not applicable.

## Data Availability Statement

The data presented in this study are available on request from the corresponding author.

## Acknowledgments

A.P. thanks the International PhD Program in Neuroscience and G.I. thanks the Ph.D. program PNRR - D.M. 117/2023, at the University of Catania.

## Conflicts of Interest

The authors declare no conflict of interest.

## Notes

### Competing Interest Statement

The authors have declared no competing interest.

